# The transcriptional dynamics of two filmy ferns from Hymenophyllaceae with different niche preferences unravel key aspects of their desiccation tolerance and vertical distribution along host trees

**DOI:** 10.1101/228866

**Authors:** Giovanni Larama, Enrique Ostria-Gallardo, Graciela Berrios, Ana Gutierrez, Ingo Ensminger, Leon A. Bravo

## Abstract

Ferns from the Hymenophyllaceae family are one of the main components of the epiphytic species diversity in the Chilean temperate rain forest. Having membranous fronds of a single layer of cells, they show a poikilohydric strategy most typical from bryophytes. Although Hymenophyllaceae species shows the ability to tolerate desiccation, there are interspecific differences in their water loss kinetic. Counter-intuitively, those species that have rapid desiccation kinetic are able to reach higher host height and tolerate higher light exposure and vapor pressure deficit. Therefore, what are the mechanisms (constitutive and/or induced) responsible of the desiccation tolerance in this fern family? As this primitive fern family is closely related with mosses, it can be hypothesized that desiccation tolerance in this particular group would be associated with constitutive features rather than induced responses during dehydration. However, the inter-specific differences in water loss and vertical distribution would be associated to different degrees of induction either within the dehydration or rehydration phases. We applied an ecophysiological transcriptomic approach to study the dynamic of gene expression in two species of filmy ferns with contrasting desiccation kinetics and vertical distribution on the host tree. Our analysis identified commonalities and differences in gene regulation, and key genes correlated with the fronds hydration state, providing the patterns of gene expression responding to microenvironmental signals and behind the physiology of their resurrection strategy

## INTRODUCTION

Among the different degrees of water deficit, desiccation is the most extreme form of dehydration affecting plants. It occurs when most of the protoplasmic water is lost and only a very small amount of tightly bound water remains in the cell matrix (Dijilianov et al., 2013). Even though evolution has shaped several plant acclimation mechanisms and adaptations to withstand short and long term periods of water deficit, plants cannot survive to desiccation, except for a unique and small group of plants called resurrection plants (Ingram and Bartels, 1996; Alpert, 2000). Desiccation tolerance is a complex trait and involves the coordination of a complex cascade of molecular events divided into constitutive and inducible molecular mechanisms to protect/repair the tissues against oxidative damage, and disruptions of metabolism and cell ultrastructure (Alpert, 2000; Ramanjulu and Bartels, 2002). The physiological and metabolic components of desiccation tolerance in resurrection plants seems to be a combination of processes underlying drought stress responses and seed maturation (Farrant and Moore, 2011; Gechev et al., 2012); however, the molecular “switches” and regulatory pathways behind desiccation tolerance is largely unknown (Dinakar and Bartels, 2013). Resurrection plants are classified into two groups according to their sensitivity and mechanisms of responses to the velocity of water loss (Oliver et al 2000). Those that can survive if water loss is rapid utilize cellular repair mechanisms, and those that survive only if water loss is gradual rely upon cellular protection mechanisms.

Resurrection plants occur in phylogenetic distinct clades and along a wide range of environments, thus the acquisition of desiccation tolerance must have occurred multiple times during plant evolution (Farrant and Moore, 2011). Therefore, the ability to survive vegetative desiccation is a widespread but at the same time uncommon in the plant kingdom (Oliver et al., 2000; Moore and Farrant, 2015). Surprisingly, resurrection plants can also be found in humid environments such as tropical and temperate rain forests (Proctor, 2003). Phillips et al. (2008) reports that *Lindernia brevidens*, an angiosperm species from the tropical rain forest exhibits desiccation tolerance. The authors suggest that *L. brevidens*, which is a species closely related with the resurrection plant model *C. plantagineum*, has retained desiccation tolerance through genome stability rather than be an adaptive response to the environmental conditions of its habitat. Another striking group of plants showing desiccation tolerance in humid habitats are the epiphytic ferns from the Hymenophyllaceae family (Pteridophyta) (Proctor, 2003). These species are called filmy ferns because they possess membranous fronds of a single layer of cells, normally lack cuticles, present no differentiated epidermis, and have no stomata (Proctor, 2012, Saldaña et al., 2013). It is suggested that Hymenophyllaceae filmy ferns are a rare example of regressive evolution from typical vascular plant adaptation to a poikilohydric strategy most typical of bryophytes (Proctor, 2012; Dubuisson et al., 2013). From an ecophysiological approach, they are adapted to live in shady and humid sites, although interspecific differences exist in niche breadth. Some species are restricted to very sheltered and high steady humidity environments such as tropical forests, while other species can inhabit sunnier sites with more marked seasonality such as temperate forests (Proctor, 2012). These geographic differences in niche breadth determine the space for the adaptive range to light and desiccation tolerance in these species. For example, species from sheltered tropical zones (e.g., *Hymenophyllum hirsutum, H. polyanthus*) show low values for light saturation points and are more sensitive to recover for mild desiccation than those species from lit sites in temperate zones (e.g., *H. sanguinolentum, H. dentatum)* (Proctor, 2012; Saldaña et al., 2013).

In the Chilean temperate rain forest, the Hymenophyllaceae family is represented by 16 species, being one of the main components of epiphytic species diversity in terms of plant morphology and habitat requirements (Parra et al., 2012, 2015). Recent studies have reported that these ferns are distributed along the whole vertical strata of their host tree; however, some species (e.g., *H. caudiculatum, H. pectinatum)* are distributed only below 60 cm where light availability is very low (10-100 μmol photons m^-2^s^-1^), while other species (e.g., *H. dentatum, H. plicatum)* extend their vertical distribution reaching heights ≥ 10 m (≥ 1000 μmol photons m^-2^s^-1^) (Saldaña et al., 2013; Parra et al., 2015). The differences in vertical distribution have been related with microenvironmental variations along host trees. Light intensity and the vapor pressure deficit increases whereas the relative humidity decrease significantly with the height of the hosts. These conditions act as a selective pressure for filmy ferns to deal with high irradiance and high evaporative demand, conditioning their microhabitat preferences (Saldaña et al., 2013). Those filmy fern species that can inhabit in the top of the host trees show better photosynthetic responses to high light irradiance in comparison to those filmy ferns inhabiting the basal strata of the host (Parra et al., 2015). In addition, the species from the top of the hosts show fast kinetics of desiccation, followed by a full physiological recovery after rehydration. Therefore, the vertical distribution pattern would be better explained by the specific microenvironmental requirements and desiccation tolerance of each species (Saldaña et al., 2013, Parra et al., 2015). This particular scenario offers a unique opportunity to address questions about the molecular and physiological mechanisms shaped by the evolutionary history of congeneric population differing in microhabitat preferences and ability to tolerate desiccation. The recent advancement in Next Generation Sequencing tools have brought important advantages to obtain large-scale datasets for the analyses of the patterns of gene expression associated to natural constraints explored in situ or under experimental conditions. Here we used RNA-seq on the Illumina Hi-seq platform to study the transcriptional responses of *Hymenophyllum caudiculatum* and *Hymenophyllum dentatum*, two Hymenophyllaceae species with contrasting vertical microhabitat preferences along host trees and different rates of water loss. Specifically, we look at the dynamics of gene expression in fronds subjected to experimental desiccation-rehydration cycles to identify commonalities and differences on gene regulation underlying the resurrection strategy associated to its water status. Our analysis identified key genes correlated with the hydration state of the fronds, and provides a broad view of the patterns of gene expression associated with the interaction of life history and their resurrection strategy.

## Materials and Methods

### Plant material, experimental design and RNA isolation.

Individuals of *H.caudiculatum* and *H. dentatum* (Fig. 1) were collected from Katalapi Park (41°31’07.5”S, 72°45’2.2”W) on winter 2014 and brought to a shaded experimental nursery garden with automated irrigation at dependencies of Universidad de La Frontera, maintaining environmental conditions as similar to the sampling site as possible. After a period of acclimation, the experimental design consisted in having fronds of both species in three different hydration states; full hydrated (FH), dehydrated (DH), and rehydrated (RH) for further comparisons of the transcriptional responses associated to each hydration state, either within and between species. The ferns went through a desiccation-rehydration process by adjusting the irrigation settings and monitoring the change in relative water content (RWC) of the fronds during seven days. For reach a fully hydrated state, ferns were subjected to irrigation pulses of three minutes at intervals of twenty minutes each, during four hours’ total; then the irrigation was shortened until reach the critical relative water content reported by the two species (see Saldaña et al., 2013), achieved at day seven (Fig. 2a). Finally, the rehydration was carried out resuming the irrigation as mentioned for the full hydrated state. The progression of desiccation was monitored by measuring the RWC, and the maximal quantum efficiency (Fv/Fm) in attached fronds (Fig. 2a). As an additional measurement of full recovery, we used an Imaging MAXI-PAM to observe the rate of recovery of the actual quantum yield (YII) at the whole frond level of both species (Fig. 2b). Fronds were collected in liquid nitrogen immediately after the measurement of RWC at the full hydrated state, at the beginning-and-end of dehydration, and at the beginning-and-end of rehydration (detailed in Fig. 2a), and stored at −80 C°. The RNA was isolated using UltraClean™ Plant RNA Isolation Kit (Mo Bio, *Carlsbad, CA, USA)* and purified with Total RNA I kit (Omega biotek, Norcross, GA, USA) according to manufacturer’s instructions. Specifically, for RNA isolation of the dehydrated and rehydrated states, we pooled samples taken at the beginning and the end of each experimental procedure, thus capturing the earliest and late transcriptional responses.

**Fig. 1.**
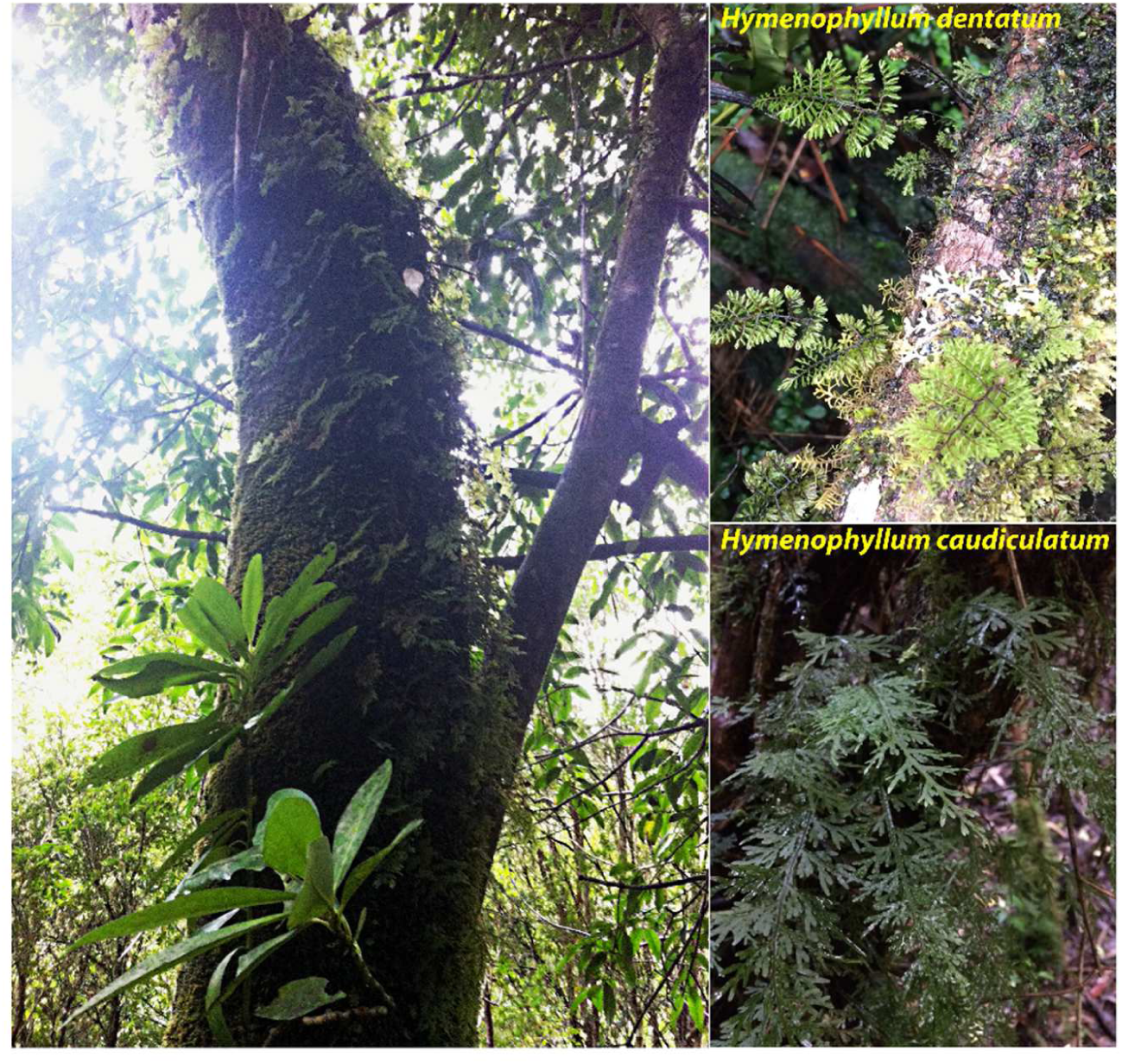
At the left of the image a reference of a host tree covered with a carpet of epiphytic filmy ferns. At the right, the two species of resurrection filmy ferns studied, attached to the trunk of their host tree.

**Fig. 2.**
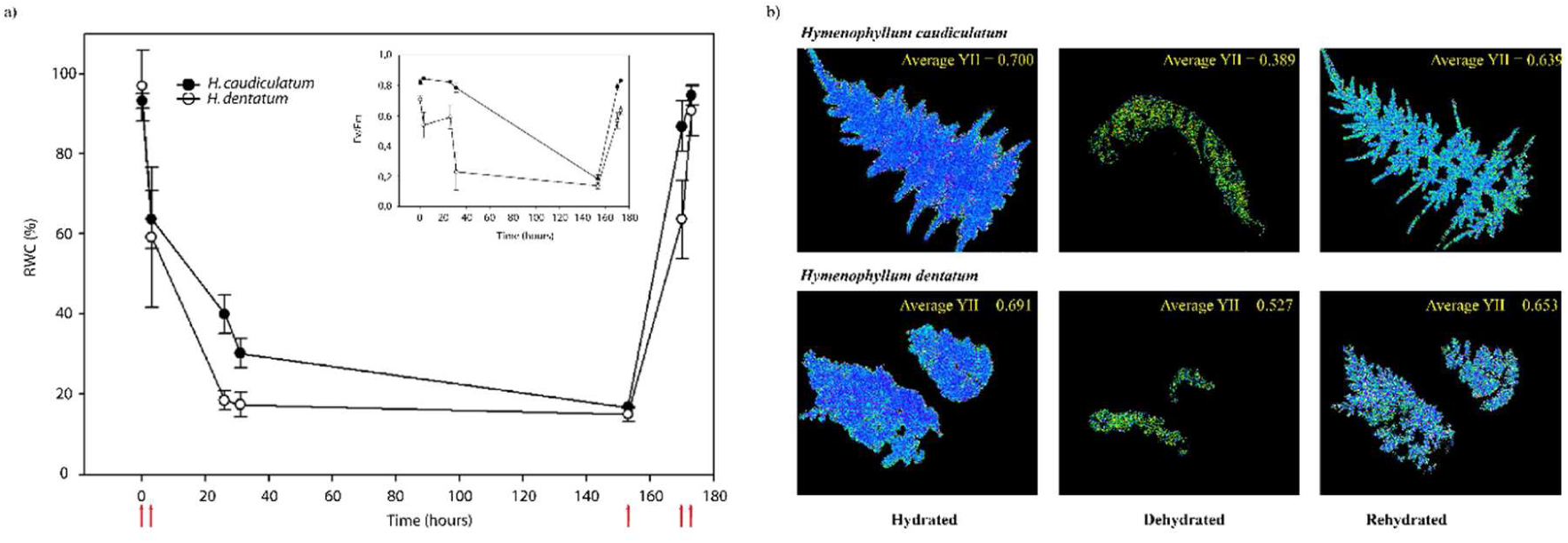
Changes in relative water content (RWC) and photosynthetic efficiency parameters (Fv/Fm, YII) of *H. caudiculatum* and *H. dentatum* during a dehydration and rehydration process (a,b). a) The two species showed a similar rate of dehydration during the first hour after irrigation shortage. After 3 hours from irrigation shortage, *H. dentatum* losses their water content faster than *H. caudiculatum.* After a week without irrigation, the two species losses approximately 80% of their water content. When the irrigation was reestablished, *H. dentatum* showed a faster rehydration rate compared to *H. caudiculatum.* Nevertheless, the maximum photochemical efficiency (Fv/Fm) was higher in *H. caudiculatum* than in *H. dentatum* (see inserted panel). Red arrows indicate sampling of fronds during the dehydration-rehydration process used for the RNAseq library construction. b) Average of the effective quantum yield of photosystem II (YII) of the entire frond of *H. caudiculatum* and *H. dentatum* measured at full hydrated, dehydrated, and rehydrated states. The two species show a decrease in YII at dehydrated state, and full recovery when rehydrated. Fluorescence images obtained by Walz Imaging-PAM fluorometer device (Walz Mess-und Regeltechnik, Germany).

### Sequencing and Illumina reads processing

The yield and quality of the RNA isolation samples was determined by an Agilent 2100 Bioanalyzer with a RIN value of six as cutoff. Then the isolated RNA was precipitated with two volumes of acetate:ethanol solution (1:10 v/v) to avoid its degradation and subsequently sent for sequencing to Macrogen Inc., Korea. A total of six samples were sequenced in a single lane of an Illumina HiSeq 2000 platform (Illumina Inc. San Diego, CA, USA) obtaining 100 bp pared end reads. The FASTQ Illumina raw sequences were processed with NGSQC Toolkit v2.3 (Patel and Jain, 2012; http://www.nipgr.res.in/ngsqctoolkit.html) to remove adaptors and low quality reads. Then, the reads were filtered based on their Q-score composition, being removed all reads with a content of Q>30 lower than 70 percent of bases (Supplemental Fig. 1).

### De novo transcriptome assemblies and refinement

The files containing high quality filtered reads for each fern were concatenated and assembled *de novo* with Trinity software package v2.1.1 (Grabherr *et al.*, 2011). Transcriptome assembly was performed at Troquil linux cluster at Centro de Modelación y Computación Científica (CMCC, Universidad de La Frontera) using 12 processors Intel Xeon E5-4640 and 192 GB of shared memory.

In order to remove misassembled and poorly supported transcripts, the reads were mapped to assembled transcripts with bowtie v1.1.1 (Langmead *et al.*, 2009), and then we estimated the relative abundance by the FPKM value (Fragments per kilobase per transcript per million mapped reads) with RSEM v1.2.26 (Li and Dewey, 2011). We kept all transcripts with ≥ 1 FPKM for downstream analysis. To handle highly similar and redundant transcripts these were clustered using CD-HIT-EST (Huang *et al.*, 2010) with a threshold of 95%.

### Functional annotation of the transcriptome

The resulting transcripts of each transcriptome were aligned into the SwissProt database using BLAST+ with an e-value filter of 1-e^-10^ as threshold, and also evaluated in hidden markov profiles to identify any family membership and conserved domains in PFAM-A database (Punta *et al.*, 2012). Functional annotation and classification was performed with PANTHER system (Mi and Thomas, 2009), using as input gene lists obtained from blast top hit using reference proteomes collection from EMBL as database. To identify any over- or under-represented groups of genes in differential expressed groups, an over-representation test was also performed using PANTHER (www.pantherdb.org).

### Differential expression analysis

The abundance estimation of the *de novo* assembled transcripts was calculated by using RSEM, which maps RNA-seq reads to the assembled transcriptomes. Reads from each sample were mapped to their corresponding final transcriptome using default RSEM parameters written in the align_and_estimate_abundance.pl script. The resulting RSEM-estimated gene abundances for each fern were merged in a matrix and analyzed with run_DE_analysis.pl script from Trinity, which involves the Bioconductor package edgeR in R statistical environment (Robinson and Oshlack, 2010; R Development Core Team, 2011). Transcripts with very low estimated counts (2 for combined groups), were not considered for edgeR pair-wise comparison of hydration states. To judge significance of gene expression, we used a False Discovery Rate value (FDR) lower than 0.05 and a minimum fold change (FC) of 2 as thresholds (see supplemental datasets 1 and 2).

### Self organizing maps (SOM) analysis

Normalized RSEM-estimated counts of both *H. caudiculatum* and *H dentatum* that fitted the expression values determined from the model described above were used for the clustering method (Chitwood *et al.*, 2013). Specifically, only genes that vary significantly in expression across fronds hydration state of both filmy ferns species were analyzed. To focus only on gene expression profile, and at the same time avoid biases with the differences in the magnitude of gene expression, the values were mean centered, selected from the upper 50% quartile of coefficient of variation, and variance scaled using the scale function (R base package; R Development Core Team, 2012) separately in *H. caudiculatum* and *H. dentatum.* The scaled expression values were used to cluster the genes of both species across fronds hydration states into a multidimensional 2 x 3 hexagonal SOM using the Kohonen package on R (Wehrens & Buydens, 2007). 100 training interactions were used during clustering with a decrease in the alpha learning rate from ca. 0.0018 to 0.0010 (Supplemental Fig. S2). After the iteration process, the final assignment of genes to the winning units form the clusters of genes (here after Nodes) associated to the hydration states. SOM outcome was visualized into pie charts for codebook vectors to obtain the counts number and mean distance of the genes assigned to each Node (Wehrens & Buydens 2007; Supplemental Fig. S3), and we used the box plot option from the ggplot2 package on R to visualize the gene accumulation patterns associated to the hydration states of the fronds in each Node. Finally, the genes of each Node were analyzed for GO enrichments terms at a 0.05 false discovery rate cutoff.

## Results

### Sequencing, *de novo* assembly and transcriptome annotation

A total of 111,495,169 and 110,988,488 paired-end reads (101 pb) were obtained after sequencing libraries of *H. caudiculatum* and *H. dentatum*, respectively, on the Illumina HiSeq2000 platform (Supplementary table 1). Following a removal of low-quality reads and duplicated reads (Supplemental Fig. S1), we performed a *de novo* transcriptomes assemblies with Trinity software by using a set of ∼85 million reads for *H. caudiculatum* and ∼87 million reads for *H. dentatum* (Supplementary Table 1). The initial assemblies resulted in 161689 contigs for *H. caudiculatum* and 332003 contigs for *H. dentatum*, which went through refinement to remove low supported transcripts. Transcripts with an estimated abundance lower than 1 FPKM and highly similar or redundant transcripts with a sequence similarity higher than 95% were removed. The resulting transcriptomes are represented by 34726 contigs for *H. caudiculatum* and 69599 contigs for *H. dentatum.* The final transcriptomes showed an increment in their N50 and average length values for both species (Supplementary table 2), indicating a removal of mainly small transcripts. Although the number of transcripts decreased significantly during the refinement, it was possible to map *c.* 80% and 70% of high quality reads to *H. caudiculatum* and *H. dentatum* transcriptomes, respectively.

Transcripts of the final transcriptomes were aligned to the SwissProt database with an alignment rate of around 50% for the transcripts of each transcriptome. In spite of the low identification rate, most of the unknown transcripts (∼80% in *H. dentatum*, and ∼65% *H. caudiculatum;* details in supplementary Table 3) belongs to small size transcripts (< 1000 bp; see Fig 3a.). An insight into the taxonomic distribution of top blast hits of transcripts revealed that both *H. caudiculatum* and *H. dentatum* had among their top hits a high amount of sequences belonging to the model moss *Physcomitrella patens* and the lycophyte *Sellaginella moellendorffii* (Fig. 3b), which is consistent with their poikilohydry strategy and the regressive evolution hypothesis (Dubuisson et al., 2013).

**Fig. 3.**
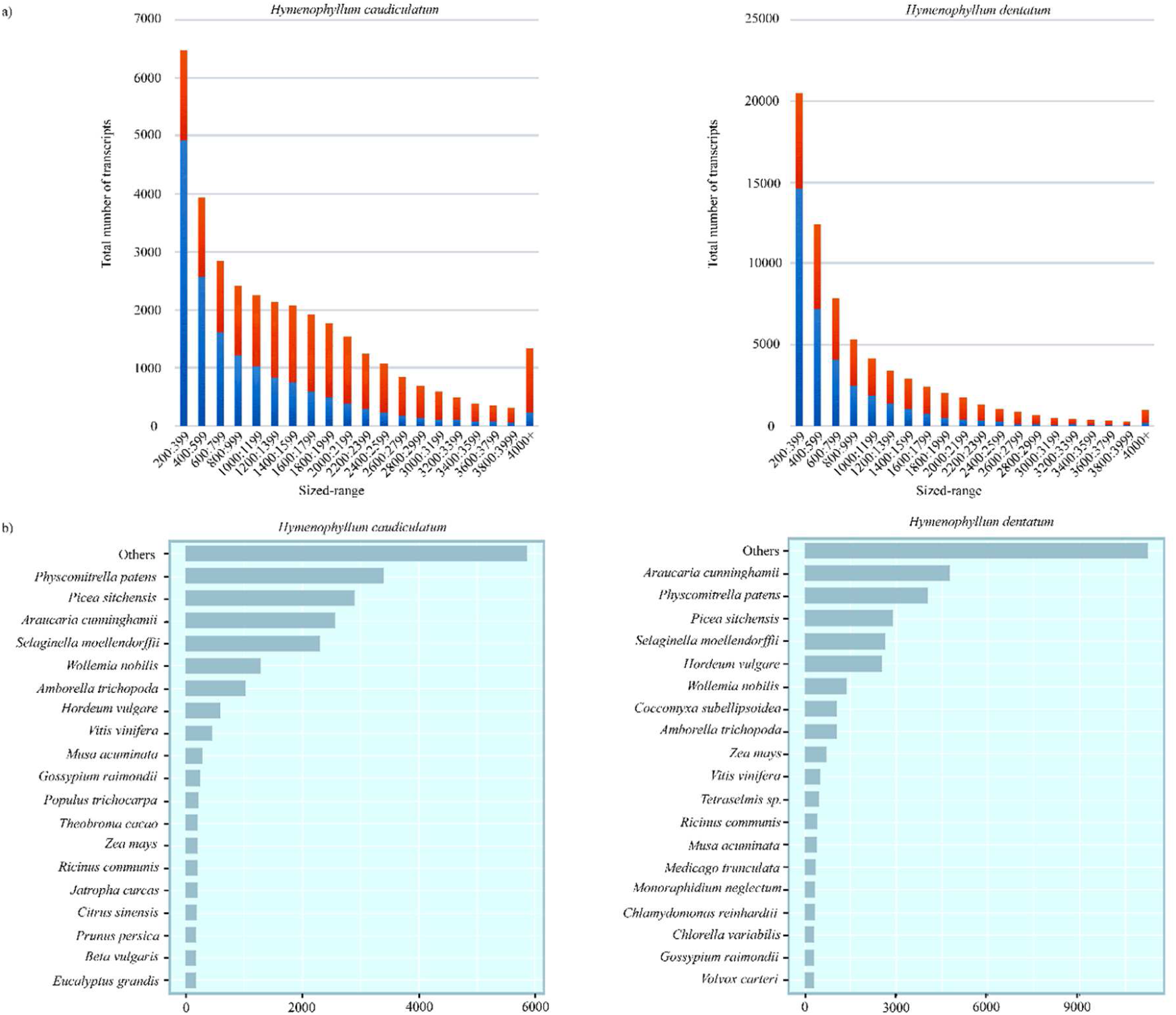
a) Transcript size distribution showing the proportion those sequences with blast hits (blue) and without blast hits (red) in the final transcriptome assemblies. b) Top-hit species distribution of the final transcriptome of *H. caudiculatum* and *H. dentatum* showing abundance of top hits to sequences of Bryophyta, Lycophyta, and Pinophyta.

### Differential expression analysis

From the final transcriptomes, we examined the expression dynamics of annotated genes during the desiccation-rehydration cycle by pairwise comparisons by using a fold change ≥ 2 and a FDR < 0.05 as cut-off (Supplemental datasets 1 and 2). For H. caudiculatum, the highest number of differentially expressed (DE) genes occurred during the dehydration process with a total of 265 DE genes, where most of them (139) were up-regulated. (Fig. 4). In *H. dentatum*, the number of up-and-down regulated genes was similar. However, among the different hydration states, rehydration presents the highest number of down-regulated genes. When comparing both species, *H. caudiculatum* presents ca. twice DE genes than *H. dentatum* and a higher proportion of up-and-down regulated genes under the dehydration process. By the other hand, *H. dentatum* showed high proportion of down-regulated genes when going through dehydration to rehydration (Fig. 4).

**Fig. 4.**
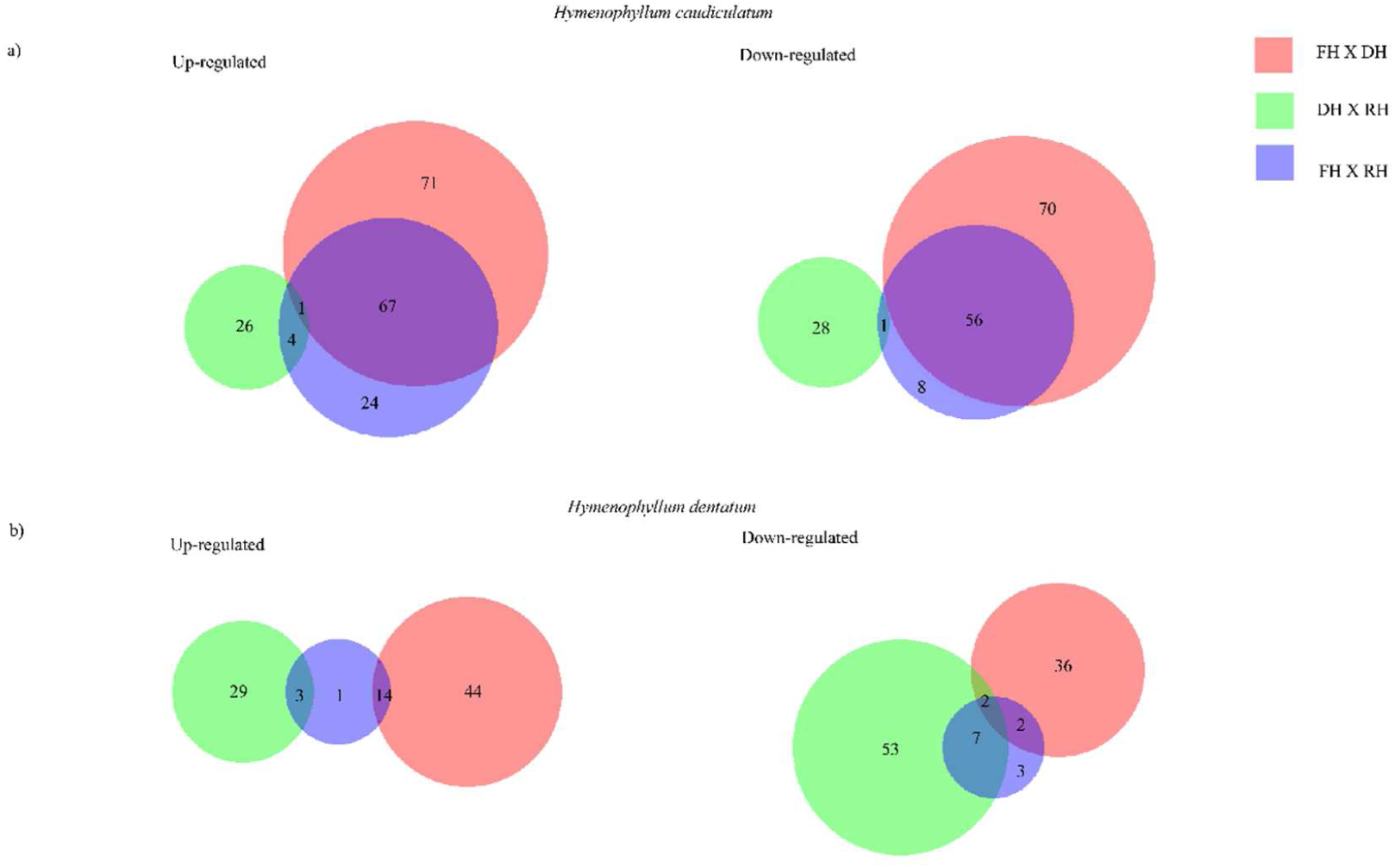
Venn diagrams showing up-and-down regulated genes in *H. caudiculatum* (a) and *H. dentatum* (b). The size of each circle is proportional to the number of unique, annotated, and non-duplicated up-or-down regulated genes in each of the species. Numbers inside the circles indicate the number of genes differentially expressed by each pair-wise comparison among the hydration states of the fronds (FH, full hydrated; DH, dehydrated; RH, rehydrated). In *H. caudiculatum*, the comparison between FH X DH contributes with the largest number of differentially expressed genes (up-and-down regulated). For *H. dentatum*, the comparison between FH X DH contributes with the highest number of up-regulated genes, whereas DH X RH contributes with the highest number of down-regulated genes. Venn diagrams were generated by using BioVenn (Hulsen *et al.*, 2008).

### GO *enrichment and expression profile of H. caudiculatum and H. dentatum*

From the differentially expressed transcripts we explored the function of the genes products by conducting a Gen Ontology analysis (GO) (See Material and Method for details). Both species showed the same enrichment pattern of sequences for each GO category (Fig. 6-must be change to 5). At the Biological Process category (BP), there was a high number of sequences in the metabolic process (> 4000 sequences) and in cellular process (> 2000). In the Cellular Component category (CC), cell part followed by organelle showed high number of sequences (ca. 1500 and 1000, respectively). At the Molecular Function category, despite interspecific differences in number, the highest accumulation of sequences was in the antioxidant activity, followed by binding (Fig. 6 -changes to 5).

Next, we explore all those differentially expressed genes in the pairwise comparisons with assigned GO annotation in order to identified those that would be candidate genes underlying the responses of desiccation tolerance within the two filmy ferns (Supplemental dataset 3 and 4). For a given hydration state, both species exhibited changes in expression of genes in common, but also in species-specific genes (Fig. 4). First, we studied the gene expression changes between hydrated vs dehydrated fronds. For *H. caudiculatum*, there was a high expression of genes encoding for subunits of ribosomal proteins, glycosylation (*U85A3*), shikimic acid metabolism (*HST*), aldo-keto reductases enzymes (*AKRs*), proteins of photosystems I and II reaction centers (e.g., *PSAA, PSBA, CP47, psbE*), carbon fixation (*RBL*), and mitochondrial uncoupling proteins (*PUMP5*). Other up-regulated genes were related with the cell signaling and chloroplasts signaling such as singlet oxygen responsible genes (*AAA*-*ATPase*), ethylene responsive transcription factors (*ERFs*), and master regulators of stress and defense responses (*transcription factor TGA6*). By the other hand, among down-regulated genes, we found genes related to carbon fixation, specifically for carbonic anhydrases (*CRSP*), oxygen evolving complex (POB), detoxification of xenobiotic and oxidative stress, group 7 of LEA, mechanical stress such as *Extensin 1*, flavonoids biosynthesis (*CHSY*) and phenolic compound metabolism (PPO). Interestingly, we found strong down-regulation of genes encoding for proteins of the light harvesting complexes I and II (e.g., *CP29, CP26*) (Table 1).

**Table 1.**
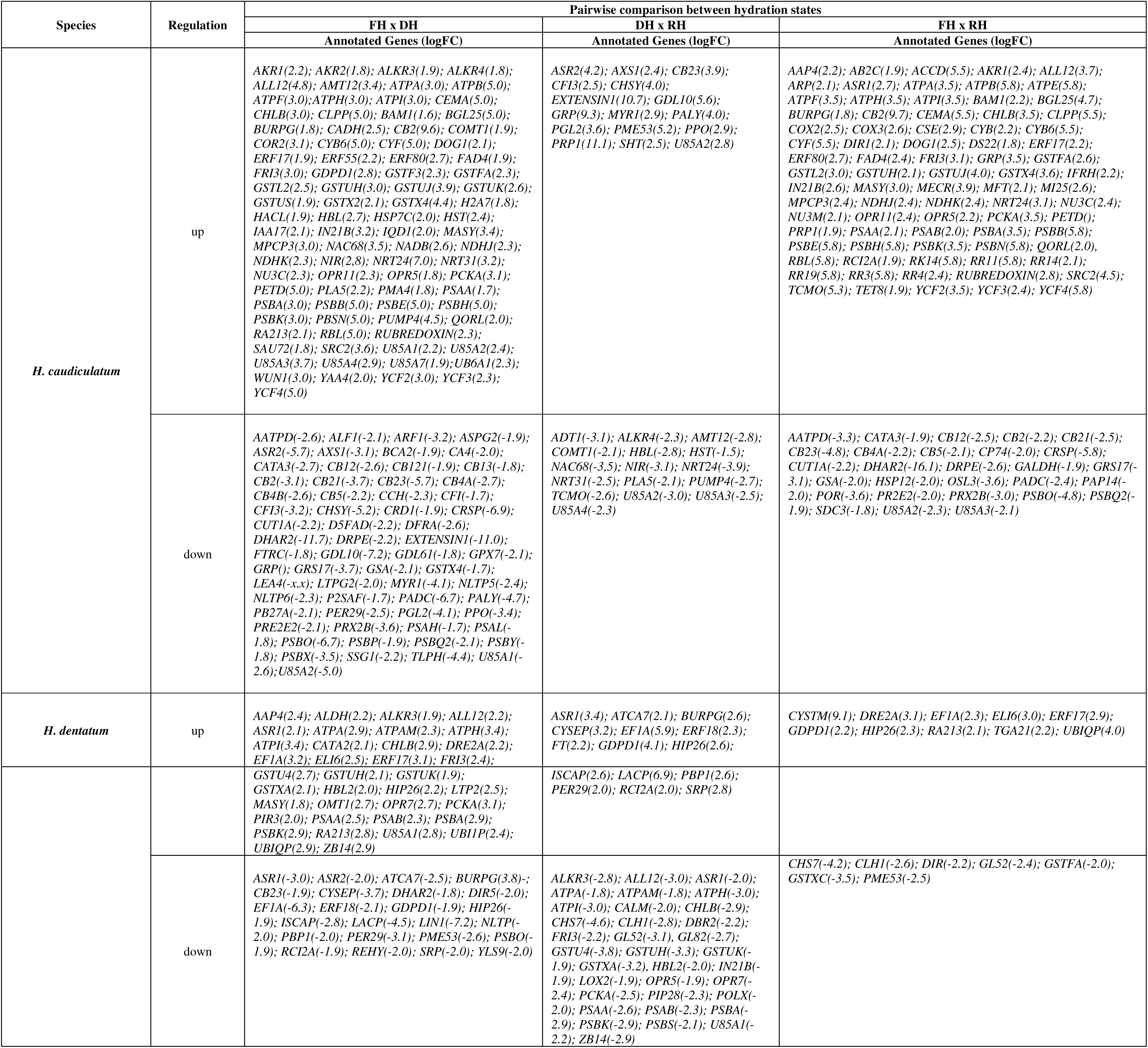
Differential expression of transcripts of interest (FDR < 0.05) during experimental desiccation-rehydration of *H. caudiculatum* and *H. dentatum* fronds. Gene identities were obtained with GO-enrichment by Panther Classification System V11.0. The level of regulation of each gene was estimated by value of the logarithm of fold change (logFC) above and below 1.0 and −1.0, for up and down regulation, respectively.

Next, during the gene expression dynamics of *H. dentatum* between hydrated vs dehydrated states, we observed up-regulation of genes related with components of chloroplast retrograde signaling (e.g., *AAA-ATPase, HIP26, OPR7, DRE2A*), genes encoding for proteins of the reaction centers of photosystems I and II, photoprotection (*early light inducible genes ELIP*), and ABA non-induced responses to stress (*oxophytodienoeate reductase OPR*). By the other hand, we found strong down-regulation of genes related with carbon fixation (*alpha carbonic anhydrase ATCA*), light harvesting complexes (*CB23*), oxygen evolving complex (*PSBO*), abscisic stress ripening protein (ASR), and protein storage (*BURP*).

Then, we studied the gene expression dynamics during the dehydration-rehydration cycle. For *H. caudiculatum*, we found upregulation of genes involved in phenolic compounds metabolism (*phenyl alanine ammonia lyase, PALY; polyphenol oxidase, PPO; phenolic acid decarboxylase, PADC*) and flavonoids biosynthesis (*chalcone synthase, CHSY*), light harvesting complexes (*CB23*), cell wall structure (*PME53*), immune system (GDSL). As down-regulated genes, we found non-symbiotic hemoglobins (*HBL*) which functions in the signal transduction pathways of several hormones, mitochondrial uncoupling protein (*PUMP4*), glycosylation (*U85A3*), aldo-keto reductases (*ALKR4*), shikimic acid metabolism (HST), nitrogen mobilization (*NIR, NRTs*). Lastly, for the gene expression changes of *H. dentatum* between dehydrated vs rehydrated states, we found upregulation of genes related with antioxidant system (*PER29*), carbon fixation (*ATCA7*), assembly of iron-sulfur proteins in chloroplast (*ISCAP*), stress responses (*SRP, ERF, RCI2A, ASR1*), detoxification mechanism (*HIP*), and chloroplast membrane lipid transformation (*GDPD*). Interestingly, we found down-regulation of genes related with structure of photosystems I and II (*PSBA, PSBS, PSAA, PSAB*), subunits of ATP synthase complex (*ATPA, ATPH, ATPI*), signaling (*HBL, CALM*), Aquaporin (*PIP28*), flavonoid metabolism (*ALL12*), Jasmonic Acid precursor (OPR7), glutathione metabolism (*GSTUK, GSTX, GSTUH*), mRNA degradation (*ZB14*), and senescence (SRG).

### Patterns of transcript accumulation across dehydration-rehydration process

A major purpose of this study was to investigate the dynamics of gene expression in two resurrection filmy ferns and to identify transcripts with similar expression patterns in response to their hydration status. The self organizing maps results (SOM) partitioned the transcripts into six clusters (hereafter Nodes) arranged as a map and, therefore, with an underlying topology showing distinct accumulation patterns of genes membership across Nodes (Fig. 5 a-c…must be changed to 6), and prominent densities of transcripts for a given hydration state within a Node (Fig. 5 b-d). Thus, those Nodes with high gene counts membership have the lowest Euclidean distance among members within the Node, which in terms of the topology, reflects the relationship between the gene coexpression in each identified Nodes.

**Fig. 5.**
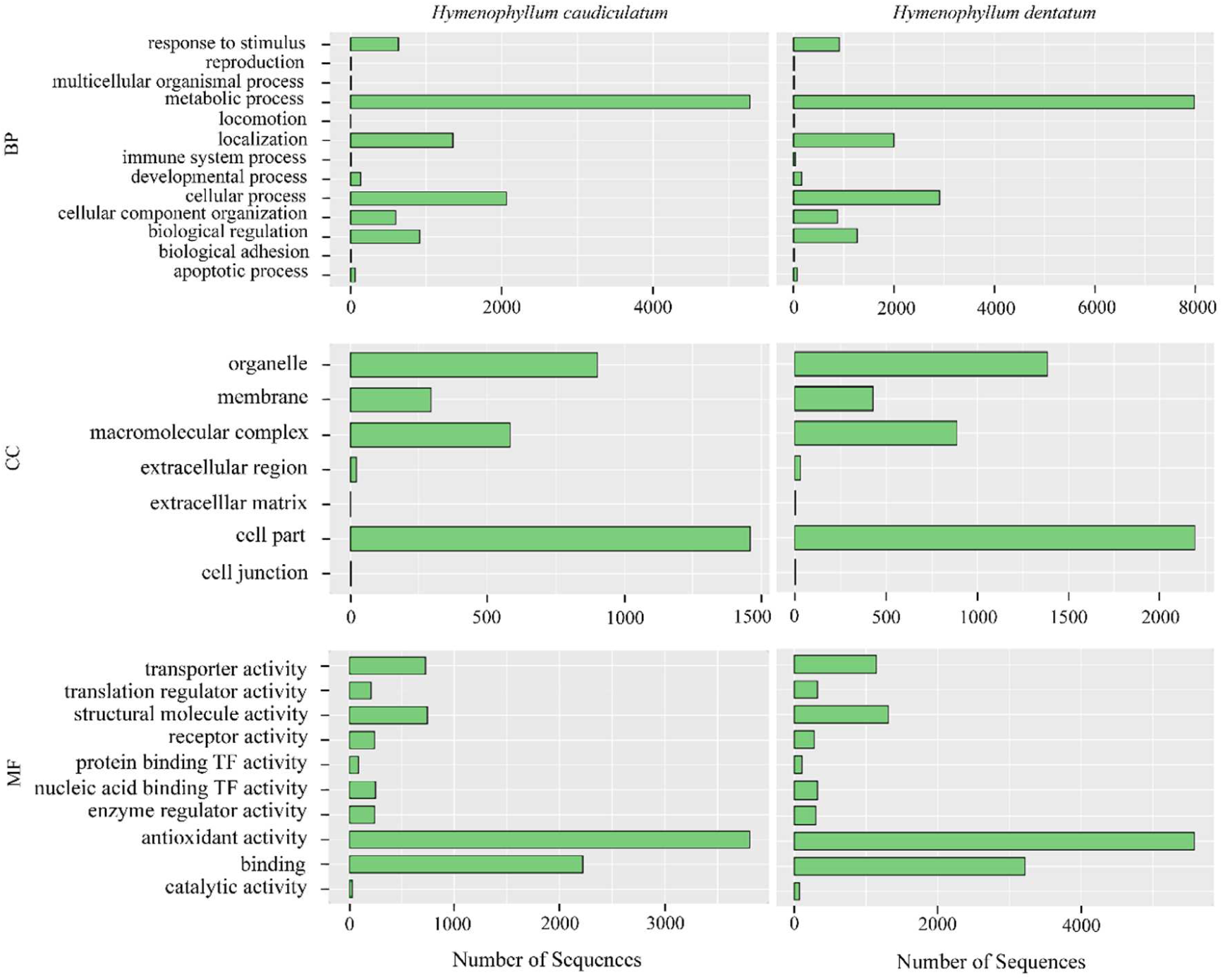
GO-category distribution of annotated genes for *H. caudiculatum* and *H. dentatum* among level 2 GO categories: biological process (BP), cellular component (CC), and molecular function (MF).

Specific Nodes exhibit high counts memberships associated to the hydration states of the ferns. For example, SOM Node 6 in *H. caudiculatum* and SOM Node 1 in *H. dentatum* have a prominent density of differentially expressed genes associated with the full hydrated state (FH). For the dehydrated and re-hydrated conditions, respectively, the enrichment of differentially expressed transcripts was observed in Nodes 2 and 5 in *H. caudiculatum* (Fig. 5b), and Nodes 2 and 6 in *H. dentatum* (Fig. 5d). In order to ascertain if SOM-based clustering yields biologically relevant results, we explored the GO enrichment of those transcripts that were enriched in particular Nodes, especially those that form prominent densities of differentially expressed genes for a given hydration state. Each Node was enriched in specific GO terms, particularly under the functional categories of biological processes (BP) and molecular function (MF)(see Fig. 6). Specifically, focusing in the gene expression changes between dehydrated vs rehydrated sates (Supplemental Dataset 1 and 2), we observed that in *H. caudiculatum*, transcripts of the dehydrated sate (Node 2) were enriched in functional categories related to stress signaling and response, photosynthesis and photosystem II stabilization and repair, unsaturation of fatty acid, and lignin biosynthetic process, whereas transcripts in the rehydration state (Node 5) were enriched in functional categories related with responses to oxidative stress, lignin biosynthetic process, photosynthesis, protein-chromophore linkage, cellular redox homeostasis, and translation. By the other hand, in *H. dentatum*, the dehydrated state (Node 2) was enriched in functional categories related mainly with antioxidant responses such as glutathione metabolic process and ROS detoxification systems, drought response, transcription, translation regulators, photosystems II stabilization, ATP synthesis and proton transport, photoprotection, ABA non-regulated stress responses. For the rehydrated state (Node 6), transcripts were enriched in functional categories corresponding mainly to ethylene and abscisic acid related signaling, photosynthesis, proton transport, plasmodesmata-mediated intercellular transport, response to stress and to toxic substance.

**Fig. 6.**
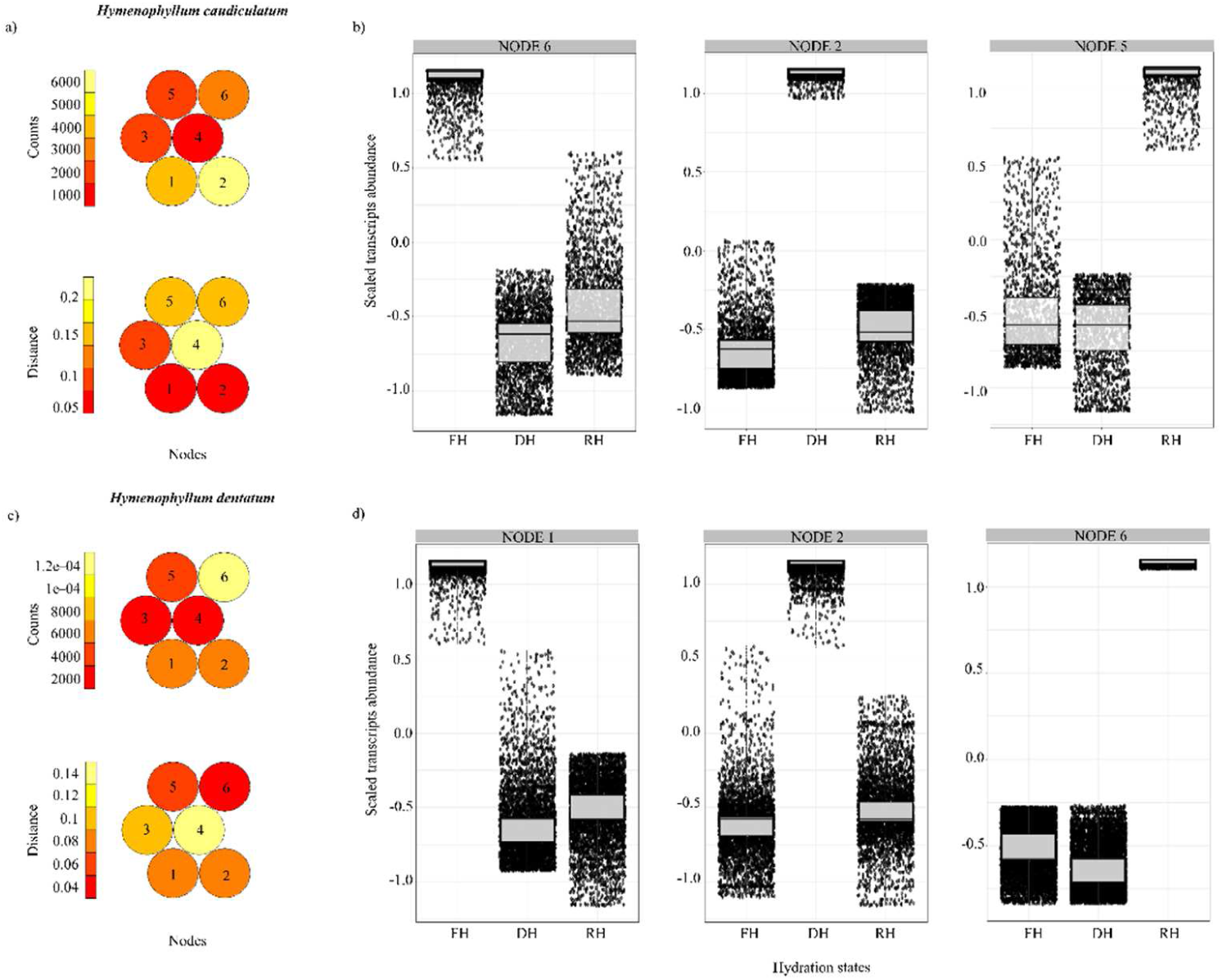
Mapping quality and clustering results for self organizing maps (SOM) of *H. caudiculatum* (a-b) and *H. dentatum* (c-d). The heatmaps shows the number of genes (counts) assigned to each SOM Node (numbered from 1 to 6) and the mean Euclidean distance of genes (distance) mapped to the particular Nodes. The constructed maps with a 2x3 hexagonal topology shows a reasonable spread out and small distances over the maps, evidencing a good mapping process. Nodes with similar codebook vectors (i.e., the patterns of gene neighboring at each hydration state) lie closer to each other. Red color indicates low count and distance, whereas light yellow indicates high count and distance. From the total six Nodes defined after SOM, boxplots were used to visualize and select those Nodes describing the highest pattern of differentially expressed genes for a given hydration state in *H. caudiculatum* (b) and *H. dentatum* (d) (see details in the Result section). For each boxplot, horizontal line represents the median, and bars represent the maximum and minimum values of the scaled gene abundance. X axis labels read as follow. FH = Full Hydration; DH = Dehydration; RH = Rehydration.

## Discussion

Because of the limited control of the fronds to control water loss, and also the patterns in habitat preferences along the host trees (Saldaña et al., 2013, Parra et al., 2015), we assumed that the resurrection strategy of these two filmy ferns depends on the sensing and responses to variations in the water status of the fronds along with the adaptation to environmental features associated to the microhabitats they occupy in their natural environment. Our study pointed to unravel the transcriptional responses of two resurrection filmy ferns to a fast dehydration-rehydration process to study the underlying mechanisms of their desiccation tolerance. Both *de novo* assembled transcriptomes (the firsts in the Hymenophyllaceae family) showed good quality parameters, comparable to other non-model species transcriptomes sequenced under the same Illumina platform and using the same criteria for subsequent downstream assembly pipelines (Xiao et al., 2011, Ranjan et al., 2014, Ostria-Gallardo et al., 2016). Thus, the robustness of our transcriptomes allowed us to explored the differential gene expression and to identify the dynamical enrichment of up-and-down regulated genes associated to the fronds hydration states, providing a comprehensive view of the response mechanisms involved in a fast dehydration-rehydration process.

Finding patterns in massive gene expression data that be consistent with a biological sense is always a challenge. Despite exploratory statistical models such hierarchical clustering (Eisen et al., 1998) and *k*-means clustering (Tavazoie et al., 1999) have undeniable advantages for identified patterns of genes with significant factor-by-species interaction effects, these methods are subjective due to human bias based on arbitrary statistical significance threshold, does not scale well or does not consider the topology between clusters (see Chitwood et al., 2013 for details of clustering methods used in multidimensional analysis). Because the characteristic of an artificial neural network, SOM results have a network topology influenced by clusters neighboring that depict biological systems (Kohonen, 1997; Wherens and Buydens, 2007). This provides an excellent tool to detect clustering patterns of gene expression across multiple factors (Wehrens et al., 2007; Clark & Ma’ayan 2011; Ranjan et al., 2015; Ostria-Gallardo et al., 2016). Our results showed prominent accumulation patterns of transcripts associated to the fronds hydration states (Fig. 5, must be changed for 6), thus we were able to elucidate both shared and species-specific molecular components associated to mechanisms of signaling and responses to desiccation (Farrant 2000; Dinakar and Bartels, 2013), disturbance of chloroplast homeostasis (Glaber et al., 2014; Chan et al., 2016), and photoinhibition related stresses (Rodriguez et al., 2010; Charuvi et al., 2015; Hayami et al., 2015). At full hydrated state, the patterns of transcripts accumulation in both species reflects the normal functioning of primary metabolic process such as photosynthesis and respiration. At dehydration, the clustering patterns and abundance of differentially expressed transcripts based on fold change allowed us to identify shared mechanisms associated with the maintenance of redox homeostasis, stabilization and repair of the photosynthetic apparatus, and chloroplast operational signaling. By the other hand, the species-specific responses were particularly associated to metabolic pathways of phenolic compounds (e.g., phenylpropanoid metabolism) and photoinhibition related processes. Both general and species-specific responses would have important roles behind their ecophysiological traits (see Saldaña et al., 2013; Parra et al., 2015; Flores_Bavestrello et al., 2016) regarding the micro-environmental conditions which they endure along its vertical distribution.

## Shared patterns of transcriptional responses during dehydration and rehydration

The patterns of differential gene expression shared by the two species (as shown in Fig. 4) would be part of conserved mechanisms regardless their habitat preferences along the host trees. During dehydration both species up-regulate genes encoding for *Gluthation-S-Transferases (GST*), and down-regulate them after rehydration. When desiccated, fronds must deal with the cytotoxic effects of reactive oxygen species associated to multiple and simultaneous stress conditions, such as water deficit and light stress (Miller et al., 2009). Given the ability of GST enzymes to inactivate a wide variety of toxic compounds (e.g., hydrophobic, electrophilic and cytotoxic substrates; see Marrs, 1996) these ferns would evolve a mechanism to finely tune the GST responses against oxidative damage during the desiccated state.

We also observed a significant up-regulation of genes encoding for ferritins during desiccation which could be a more specific mechanism coupled with the aforementioned for GST. Ferritin are a putative iron storage protein that in plants have been linked with ROS metabolism. Specifically, ferritin synthesis is activated at the transcriptional level by iron and H2O2 as well as by high light intensity, enhancing ROS detoxifying enzymes (Briat et al., 2010). The evidence that link ferritin and ROS metabolism points out that the control of ferritin synthesis is required for a proper maintenance of the redox status of fronds cells. Moreover, it is known that ferritin domains are part of the structure of desiccation related proteins (DRP, see Candotto et al., 2016), therefore besides the antioxidant role of ferritin, these ferns may use this proteins for protection of membrane structures, such as photosynthetic structures (Bartels et al., 1992), because photosynthesis must be reestablished as soon as possible upon rehydration. An efficient detoxification and antioxidant defense systems are key components for desiccation tolerance (Ramanjulu and Bartels, 2002; Dinakar and Bartels, 2013; Charuvi et al., 2015), thus our studied ferns showed to be well equipped with genes involved in a highly efficient molecular control for scavenging ROS when desiccated.

Both species also shared the pattern of gene regulation which controls the structure and function of photosystems. Aside from the regulation of protein translation or degradation, the transcriptional control of photosynthetic proteins is key for the balance of chloroplast redox state (Dinakar & Bartels 2013). During desiccation both species up-regulate genes encoding for reaction centers of Photosystem I and II whereas those genes encoding for the light-harvesting antenna and oxygen-evolving complexes were down-regulated. Interestingly, this response differs with what has been reported for the model resurrection plant *Craterostigma pumilum*, in which the inverse pattern occurs (Charuvi et al., 2015). A plausible explanation for our results is that because of their homoichlorophyllous strategy (Flores-Bavestrello et al., 2016) and the short period of time it takes during dehydration and rehydration to reach the critical relative water contents (RWC) and then recover from that condition (ca. 2 hours total; see Saldaña et al., 2013), these resurrection ferns must minimize the potential for light capture and the oxidation of water in order to decrease redox reactions but maintain a stock for reaction centers. This strategy would help during the time-lapse of desiccation-rehydration against the combined effects of severe dehydration and intermittent occurrences of high irradiances inputs by sunflecks to deal with reactive oxygen species and oxidative damage and for maintain the balance of photoprotection and photoinhibition (Müller et al., 2001).

The coordination of the complex mechanisms such as those mentioned above require intertwined signaling pathways for systemic responses against the internal and external constrains associated to the resurrection strategy. The transcriptional dynamics of the studied resurrection filmy ferns showed key components of the core module involved in the chloroplast retrograde signaling (Chan et al., 2016). For example, we observed up-regulation of *AAA-ATPase* genes which are known to be involved in ROS signaling and response; also components of signal transduction such as Auxin and calcium signaling, oxylipins metabolism, Ethylene and ABA responsive transcription factors. Because the fronds of these ferns ranges among one to a few cell layers, and given that plastid-to-nucleus signaling is essential for the coordination and adjustment of cellular responses to external and internal cues (Yamaguchi & Shinozaki 2006; Glaber et al., 2014; Lepetit & Dietzel 2015), environmental perturbations imposing restrictions for the chloroplast homeostasis would have a great impact in whole frond form and function. Fluctuations in fronds water status will initiate an operational signaling to change the expression of thousands of genes to adjust chloroplast and the whole cell functioning (Chan et al., 2016). According to our analyses, the operational response triggered by desiccation would activate mechanisms to cope with mechanical, physical and biochemical related stresses. For example, both species up-regulate genes encoding for AP2/ERF transcription factors during desiccation (e.g., *DREB, ERF, RAP*), which are known to be involved in responses to environmental stimuli, especially for redox, osmotic and drought stress (Glaber et al., 2014; Chan et al., 2016). Moreover, Vogel et al. (2014) points out that the coordination of chloroplastic and nuclear gene expression is mediated by the interactions of sugar levels, redox stat that activate protein kinase phosphorylation cascades and AP2/ERF transcription factors in response to high light. Our results indicate that this would be a key mechanism used by the studied filmy ferns contributing to its resurrection strategy.

## Species-specific patterns of transcriptional responses during dehydration and rehydration

Due to the different niche preferences of both species, one of the most ambitious goals of our study was to identify species-specific transcriptional responses in order to have insights into particular physiological conditions that may prevail under different frond hydration states driven by its evolutive history. As an overview, the specific responses in *H. caudiculatum* underpins to enhance osmotic responses and phenylpropanoid related pathways, whereas in *H. dentatum*, the specific responses point to enhance protection against oxidative damage and high light stress. Examining the responses of *H. caudiculatum* during dehydration, we observed up-regulation of *DOG1* gene. The expression of this gene is known for controlling the timing of seed dormancy in responses to environmental signals such as temperature by altering gibberellin metabolism (Graeber et al., 2012). Given that during embryo development and maturation the acquisition of desiccation tolerance is a key process for the viability of the majority of seeds, it is expectable that resurrection plants use part of the molecular components involved in the control of seed dormancy. However, the fact that *DOG1* gene was up-regulated only in *H. caudiculatum* suggest a mechanism linked to ABA and GA balance by a common intermediary such as DELLAs proteins (Kendall et al 2011).

We also observed up-regulation of genes encoding degradation of starch and glycosidic bonds (Table 1). Given that *H. caudiculatum* inhabit in most humid microhabitat, dehydration imposes the major constrain to its homeostasis especially due to osmotic stress. The up-regulation and further expression of genes encoding enzymes such as β-amylase and β-glucosidase would rise the release of sugar and sugar-related osmolytes from starch degradation and improves osmotic stress tolerance (Thalmann et al., 2016). In addition, sugar-related osmolytes can also have protective effects on chlorophyll membranes and protein stability of resurrection plants (Mancini et al., 2012; ElSayed et al., 2014).

Another specifics response in *H. caudiculatum* was the significant up-regulation of genes encoding for enzymes involved in phenylpropanoids biosynthesis, specifically polyphenols. As precursor of cell wall components, this response may be linked to drought and osmotic stress for preserving the structural integrity of the cell wall during the reversible mechanism of cell wall folding and shrinking during dehydration/rehydration is a key feature for the resurrection strategy (Giarola et al., 2016). These specific responses observed in *H. caudiculatum* favoring mechanisms against drought and osmotic stress tolerance are in agreement with the reported habitat preferences (i.e., base of the trunks) for this species along the host trees. Interestingly, during rehydration this species showed high up-regulation of enzymes involved in the catabolism of phenolic compounds such as the phenolic acid decarboxylase and polyphenol oxidase which indicates an active regulation over these secondary metabolites during short but frequent events of desiccation-rehydration.

On the other hand, the specific differential gene expression responses of *H. dentatum* underpins to a physiological strategy to cope with high light stress. In particular, we observe significant up-regulation of the *ELI6* gene which encode for an early light inducible protein (Table 1). It is known that *ELIP* genes increase their expression in the responses to stress related to high light and photoinhibition, and are involved in maintaining the structural integrity of photosystems (Bartels et al., 1992; Gechev et al., 2012; Hayami et al., 2015). The up-regulation of *ELIP* only in *H. dentatum* would be a response associated with its widest distribution toward more illuminated micro-sites along the host trees. The other specifics responses are involved in respond to stressful conditions and scavenging of specific ROS such as hydrogen peroxide.

## Conclusion

Only few transcripts of the total transcriptomes dataset showed significant changes in their expression (see detailed numbers in the Result section), indicating that the mechanism of desiccation tolerance of these ferns is achieved by constitutive mechanisms, which provide a background protection against their rapid loss of water. However, those genes subjected to a highly dynamic expression that depends on the water status of the frond would be a response of inducible mechanisms triggered by signals of their particular niches preferences. The quality of our *de novo* transcriptomes assemblies allowed us to identify key genes involved in shared and species-specific responses of the studied ferns during a dehydration-rehydration process. The analysis and comparison of the transcriptional profiles these two filmy ferns under a fast dehydration-rehydration process of shed new lights into components and mechanisms of the signaling and responses involved in its resurrection strategy, and allowed to identify species specific induced mechanisms that would contribute to their ecophysiological traits and microhabitat differences in their natural environment. Finally, much more work is needed to understand the roles and contribution of specific molecules and postransciptional regulations that may have central roles in the desiccation tolerance strategy of these species when subjected to rapid loss-and-gain of water as occurs in the natural environment. Also, studies under an evolutionary developmental biology approach would help to decipher and understand the genetic architecture and the possible network rewiring associated to the microclimatic conditions in which these resurrection filmy ferns have evolved.

